# Nicotine Ameliorates *α*-synuclein Pre-formed Fibril-Induced Behavioral Deficits and Pathological Features in Mice

**DOI:** 10.1101/2024.05.09.593280

**Authors:** Zhangqiong Huang, Yue Pan, Kaili Ma, Haiyu Luo, Qinglan Zong, Zhengcun Wu, Zhouhai Zhu, Ying Guan

## Abstract

**Background:** Epidemiologic study suggests nicotine reduces risk of PD, could be potential treatment for Parkinson’s disease.

**Objective:** To study the effect of nicotine on behavioral phenotypes and pathological characteristics of mice induced by human alpha-synuclein preformed fibers (α-syn-PFF).

**Methods:** Mice were injected with 5 μg of human α-syn-PFF in the hippocampus while administering nicotine-containing drinking water (200μg/mL). After 1 month, the motor ability, mood, spatial learning, and memory ability of the Parkinson’s disease(PD)phenotype-like model were detected using open field, rotarod, Y maze, and O maze tests. The expression of pathological α-syn, apoptotic proteins and the numbers of glial cells and neural stem cells in the hippocampus of mice were detected using western blotting and immunofluorescence.

**Results:** Nicotine significantly reduced pathological α-syn accumulation, α-syn serine 129 phosphorylation and cell death caused by PFF injection in the hippocampus of mice, inhibited the increase of glial, microglia and apoptotic cells, decreased the expression levels of PI3K and Akt.

**Conclusions:** Nicotine may have inhibitory effects on human α-syn-PFF-induced neuroinflammation and apoptosis. Thus, it reduces human α-syn-PFF-induced behavioral deficits and pathological changes in mice.

## Introduction

Neuronal inclusion bodies composed of alpha-synuclein (α-syn) or Lewy bodies are a typical pathological feature in patients with Parkinson’s disease (PD) and dementia with Lewy bodies (DLB)(Spillantini et al. 1997; Baba et al. 1998). The α-syn misfolded protein aggregates to form amyloid fibrils, many of which form inclusion bodies in the cells. In recent years, it has been generally accepted that abnormal aggregation and pathological propagation of α-syn are central mechanisms in the pathogenesis of PD (Wong and Krainc 2017). It has been shown that in vitro recombinant α-syn-PFF can spread between neuronal cells like prions, Moreover, following the injection of α-syn-PFF into the brains of mice or after using it to treat cells, the S129 sites of α-syn-PFF can be phosphorylated to form p-α-syn (a hallmark of pathological α-syn) and promote the formation of insoluble α-syn from water-soluble α-syn, both of which are characteristic of pathological α-syn (Loria et al. 2017; Kam et al. 2018). Some investigators have stereotaxically injected α-syn-PFF into the striatum, substantia nigra, and olfactory cortical layers of rats or mice. The α-syn-positive inclusion bodies appear not only at the original injection site but also in other brain regions that are far from the stereotaxic injection site (Masuda-Suzukake et al. 2014; Mezias et al. 2020). Importantly, unlike viral vector-based models, α-syn-PFF models show a pathology resulting from PD-related pathological changes in endogenous α-syn after α-syn-PFF injection (Volpicelli-Daley et al. 2011; Luk et al. 2012; Paumier et al. 2015), and the resulting neuronal dysfunction is more similar to human disease (Kurowska et al. 2016). Additionally, compared to the α-syn transgenic model of PD, the α-syn-PFF models show a longer time course of degeneration, with early α-syn pathology in PD-associated brain regions and the development of dopamine dysfunction, nigrostriatal degeneration, and motor deficits several months after induction (Luk et al. 2012; Luk et al. 2012; Paumier et al. 2015; Visanji et al. 2016). Thus, the α-syn-PFF PD model, which mimics the prion-like properties of extracellular α-syn spreading between neurons, represents a novel animal model that is suitable for studying the pathological mechanisms of PD (Visanji et al. 2016; Loria et al. 2017).

Epidemiological studies have shown that substances such as caffeine, inosine, calcium channel blockers, and nicotine all reduce the risk of PD. Moreover, nicotine and caffeine are thought to affect the aggregation of alpha-syn (Oertel and Schulz 2016) and may represent new drugs for PD. Nicotine is a neuroactive compound involved in the development and maintenance of tobacco addiction (Cohen et al. 2015a; Vazquez-Sanroman et al. 2017) and is also an agonist of nicotinic acetylcholine receptors (nAChRs). Although nicotine abuse has adverse effects on brain plasticity (Collo et al. 2013; Carboni et al. 2018), in heavy smokers, nicotine abstinence is accompanied by cognitive impairment, and its acute effects on the adult brain are thought to be beneficial (Pistillo et al. 2015), particularly the neuroprotective effects. The alkaloids contained in tobacco increase the lag time of the nucleation step and reduce the accumulation of the more toxic oligomers in a concentration-dependent manner. Moreover, some researchers have found that nicotine directly inhibits the formation of α-syn fibrils, destabilizing pre-formed α-syn fibrils and increasing the stability of soluble oligomeric α-syn (Ono et al. 2007; Hong et al. 2009). Using in vitro and yeast models of PD, Kardani et al. also found that nicotine administration resulted in the inhibition of α-syn oligomer formation (Kardani et al. 2017). Clinical studies have shown that when patients with PD are given nicotine and levodopa in combination, nicotine treatment could substantially reduce the dosage of levodopa with the same efficacy while also alleviating side effects such as dyskinesia (Domino et al. 1999; Huang et al. 2011a). In addition, numerous animal experiments have shown that nicotine activates nAChRs at the terminals of dopaminergic neurons, regulates dopamine release, and prevents significant loss of striatal dopamine, thereby having a protective effect on dopaminergic neurons (Costa et al. 2001; Quik and Kulak 2002; Quik et al. 2012; Kyaw et al. 2013; Barreto et al. 2014). Recently, nicotine treatment of a *Drosophila* model of PD inhibited the Parkinson’s-like phenotype by increasing the inhibition of tyrosine hydroxylase and the level of dopamine (Carvajal-Oliveros et al. 2021).

Among patients with PD, 80% go on to develop dementia, showing cognitive impairment. The contribution of abundant Lewy neurites in the hippocampal region to dementia was understood long before the discovery that α-syn was a major component of DLB pathology (Dickson et al. 1991). The presence of α-syn inclusion bodies in brainstem nuclei is important for motor function, and their abundant presence in cortical and hippocampal regions is also predicted to be associated with the development of cognitive impairment (Tsuboi et al. 2007; Halliday et al. 2008; Irwin et al. 2017). Moreover, clinical studies have confirmed that cognitive decline in PD and Lewy body dementia is attributable to damage to the hippocampus and cortical circuits, with significant hippocampal atrophy observed in patients with PD (Calabresi et al. 2013; Braskie and Thompson 2014; Fraser et al. 2015; Pini et al. 2016). Previous studies conducted by our team have shown that overexpression of α-syn in the hippocampal region can impair neuronal survival in adult mice (Du T et al. 2021). Moreover, cholinergic input to the hippocampus affects adult hippocampal neurogenesis. Emerging neurons express functional ionotropic nAChRs during the differentiation and maturation stages, and their expression is essential for the normal survival and integration of newborn neurons into hippocampal circuits (Otto and Yakel 2019). Additionally, there is evidence that nAChRs play an important role in the differentiation, maturation, survival, and integration of newborn neurons (Campbell et al. 2010). In contrast, chronic nicotine exposure affects various developmental stages of newborn neurons, while experimental delivery of high doses of nicotine decreases the proliferation, differentiation, and survival of newborn neurons (Wei et al. 2012). Long-term oral administration of low-dose nicotine to a transgenic mouse model overexpressing α-syn demonstrated that nicotine significantly improved cognitive deficits induced by α-syn overexpression but did not alter α-syn aggregation(Subramaniam et al. 2018). So, more evidence is needed regarding nicotine’s ability to ameliorate behavioral deficits by inhibiting α-syn aggregation, astrocyte activation, and neuronal apoptosis.

In this study, we established a PD phenotype-like model by injecting recombinant human α-synuclein pre-formed fibril (α-syn-PFF) into the hippocampal region of mice, while drinking nicotine-containing water, and then we examined the motor function, cognitive function, mood changes, and pathological changes occurring in the hippocampal region of the mouse model. To investigate the effect of nicotine on the behavioral deficits induced by α-syn-PFF in mice, and to further investigate the molecular mechanisms, and to provide the basic materials for the study of nicotine amelioration of the hippocampal neuronal damage caused by α-syn-PFF.

## Materials and methods

### Animal

Forty male C57BL/6 mice aged 6–8 weeks and weighing 20–24 g were purchased from the Institute of Medical Biology, Chinese Academy of Medical Science, Kunming, China) and housed in a barrier environment maintained at a constant temperature (20–24°C) and relative humidity of 50–60%. The mice had free access to water and food, and the cages were under a 12/12-h day/night cycle, with regular twice-weekly cleaning. All mice were allowed to acclimatize to the housing environment for 1 week before the start of the experiment. The experiment was approved by the Experimental Animal Ethics Committee of the Institute of Medical Biology Chinese Academy of Medical Sciences (DWSP202203024), and the experimental procedure followed the 3R principle of giving humane care to the animals.

### Preparation of recombinant human *α*-syn-PFF

The highly efficient prokaryotic expression vector pGEX-5X-1-syn was constructed and preserved in our laboratory and was induced by 1.0 mM IPTG at 37°C for 3 h after the transformation of the BL21 strain (preserved in our laboratory). Following the isolation of the supernatant, the protein was purified using high-efficiency GST fusion protein purification magnetic beads (Beads Beaver Biosciences Inc., China).

After purification and concentration, 9 mg/mL of human α-syn was incubated for 6 d at 37°C with 1000 r/min shaking. After the preparation was verified using western blotting and Coomassie blue staining, prepared human α-syn-PFFs were diluted to 2 mg/mL, divided among 50-μL tubes, and stored at −80°C. Subsequently, α-syn-PFF was sonicated with a water-bath ultrasonic crusher for 1 h at a low temperature before stereotaxic surgery.

### Establishment of a mouse model of human *α*-syn-PFF

Mice were treated with a single bilateral injection into the dorsal hippocampus. The mice were preoperatively injected intraperitoneally with a ready-to-use anesthetic containing 1.25% avertin at a dose of 0.2 mL/10 g. The coordinates of the injection referred to the Atlas of Mouse Brain Stereotaxic Positioning: anteroposterior (AP) −2.5 mm, mediolateral (ML) ± 2.0 mm, and dorsoventral (DV) −2.0 mm. After placing the mice under anesthesia, the scalp was incised for hippocampal localization drilling, and 2.5 μL of α-syn-PFF was injected into both hippocampi using a 5-μL Hamilton microinjector, while the control group was injected with sterile phosphate-buffered saline (PBS). After injection, the needle was left in place for 5 min and then slowly removed and the scalp was sutured. After surgery, the mice were placed in a thermostatic recovery apparatus and fed routinely after they had woken. Behavioral tests were performed after 1 month of feeding.

### Grouping of mice and nicotine treatment

Forty mice with similar locomotor ability were randomly divided into a normal control (CON) group, an α-syn-PFF group, an α-syn-PFF + nicotine group, and a nicotine group. (i) The CON group mice were injected with sterile PBS in both hippocampi and fed normal drinking water. (ii) In the α-syn-PFF group, mice were injected with human α-syn PFF in both hippocampi and fed normal drinking water. (iii) In the α-syn-PFF + nicotine group, mice were injected with human α-syn-PFF in both hippocampi and fed drinking water containing 200 μg/mL nicotine. (iv) In the nicotine group, mice were injected with sterile PBS in both hippocampi and fed drinking water containing 200 μg/mL nicotine.

### Behavioral test

#### Open field test

The mice were allowed to acclimatize to the test environment for at least 2 h before the test. A 50 × 50 cm field was divided into two areas, the center and the periphery, with the sides of the center being approximately half the length of the periphery. The mice were placed at a uniform distance from the center and observed for 5 min to test their spontaneous activity.

#### Rotarod test

We tested the spontaneous activity of mice in a spacious environment. The total test time was 5 min, where the first 2 min were set for accelerated exercise to test the coordination of the mouse’s limbs, and the last 3 min were set for uniform exercise to test the mouse’s physical strength. The maximum speed was 40 rpm/h. Each mouse was tested three times at least 3 min apart, and the time taken for the mouse to fall off the spinning bar was recorded.

#### Y maze test

According to a previously published method (Kraeuter et al. 2019), a Y maze test was used to assess the short-term memory of mice. During the training period, one arm of the Y maze (the novel arm) was closed, and the mice were placed in the maze from the starting arm and allowed to explore freely for 10 min. After an interval of 1 h, the novel arm was opened, and the mice were re-placed and allowed to explore the maze freely for 5 min. The number of entries into the new arm was recorded and compared with the number of entries into the other arm to assess the degree of spatial memory.

#### O maze test

The anxiety-like behavior of mice was examined using the rodent’s preference for dark enclosed spaces. The mice were placed in an O maze for 5 min, and the number of times they entered the open arm and the time they spent in the open arm were recorded.

### Western blot assay

The brains of mice were obtained after sacrifice, and the hippocampus was separated and placed in a cold eppendorf tube. Subsequently, the protein samples were separated with RIPA lysate buffer (CW2333, CWBIO, and China) containing 2 mM Phenylmethanesulfonyl fluoride (PMSF) and protease inhibitor mixture (539,131, Millipore, and USA). The supernatant obtained after centrifugation was taken for protein quantification using a BCA kit (CW0014S, CEBIO, and China). Subsequently, 5× loading buffer was added, and the sample was boiled for 5 min. Isolation was performed on Criterion TGX non-staining gel at 85 V for 120 min. After transfer to a semi-dry transmembrane, 5% skim milk was added at room temperature for 45 min. To avoid wasting resources and increase efficiency, we then cut the membrane with the blots. The membranes were incubated with primary antibody at 4°C overnight, before adding secondary sheep anti-mouse at room temperature for 1 h. Odyssey software was used for quantification.

### Immunofluorescence

Mice were anesthetized with 1.25% avertin and then perfused with PBS and 4% paraformaldehyde solution. The brains were fixed in 4% paraformaldehyde solution overnight, dehydrated, and sliced into 4-μm-thick sections after embedding. The slices were oven dried at 65°C, routinely dewaxed and hydrated, microwave boiled in Tris-EDTA buffer for 15 min for antigen repair, cooled to room temperature, and then incubated with goat serum for 1 h. The slices were incubated overnight at 4°C with the corresponding diluted primary antibody, rinsed in PBS, and then treated with a fluorescent rabbit secondary antibody. The images were captured using Panoramic MIDI (3DHistech, Hungary) after DAPI (Abcam, ab104139) blocking.

### TUNEL staining

Hippocampal cell apoptosis was detected using YF®594-dUTP notch nick-end labeling (TUNEL) mediated by terminal deoxynucleotide transferase. The test procedures were performed according to the manufacturer’s instructions (YF ® 594 TUNEL Assay Apoptosis Detection Kit; US Everbright Inc., China, T6014). The nuclei were stained with DAPI (Abcam, ab104139).

### Statistical analysis

All behavioral images were recorded using the SMART V3.0 behavioral video analysis system. GraphPad Prism 8.0 (GraphPad Software Inc., San Diego, CA, USA) was used for statistical analysis. Kruskal-Wallis was performed for multiple-group comparisons. The images were processed using Adobe Photoshop CS6. All data are shown as the mean ± standard deviation (SD). P-values < 0.05 was considered significant (“*” means P < 0.05; “**” means P < 0.01; and “***” means P < 0.001).

## Results

### Testing of the mouse model constructed by hippocampal injection of *α*-syn-PFF and detection of nicotine intake

Western blotting and Coomassie blue staining were performed on the prepared α-syn-PFF with antibodies that specifically recognized human α-syn. A large amount of macromolecular protein was detected in α-syn fibrils(45kDa) (Fig. 1A,B), while a long fiber structure was observed under a transmission electron microscope (Fig. 1C). The above results confirmed the successful preparation of recombinant α-syn fibrils.

**Fig. 1.**
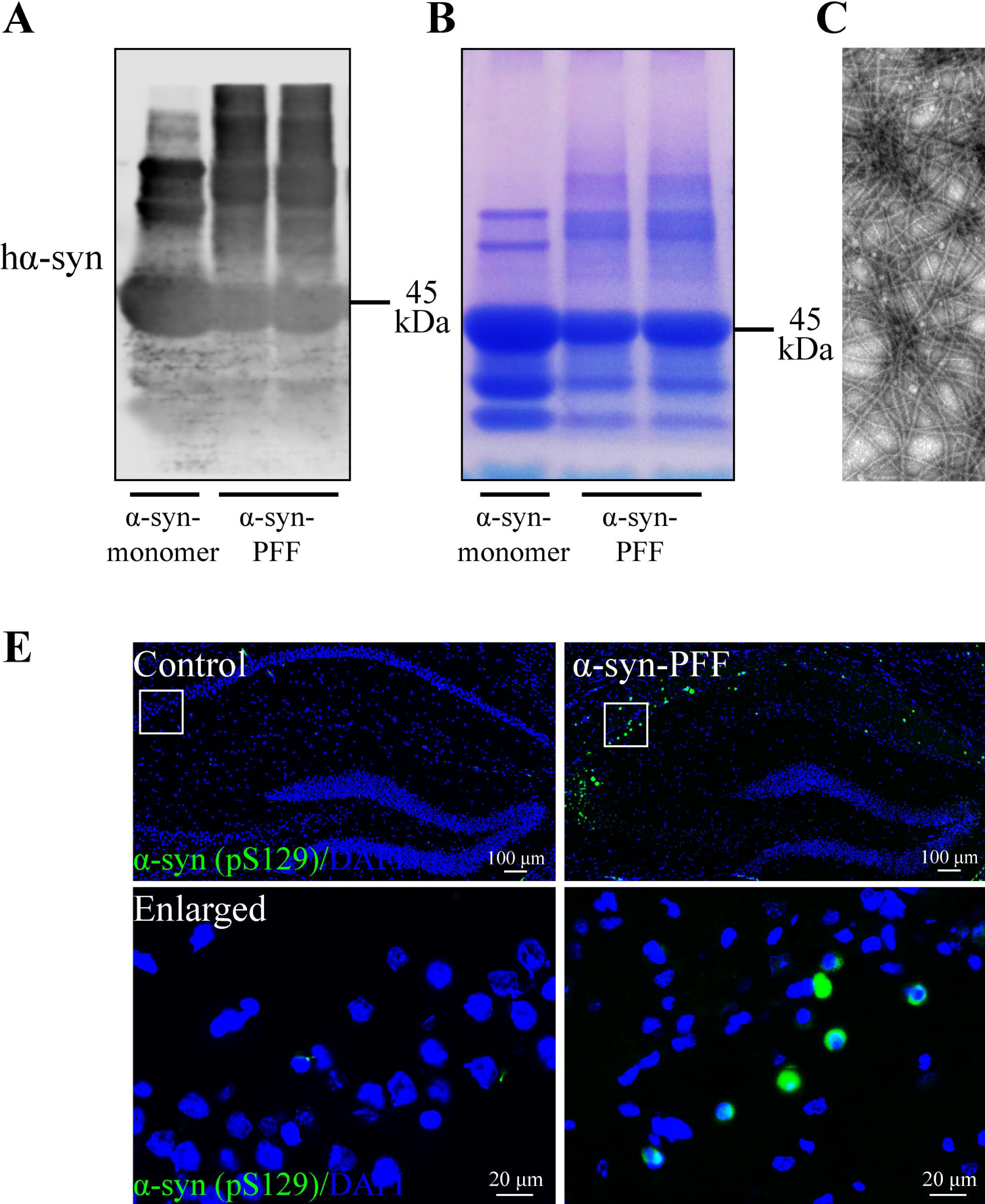
Nicotine inhibits α-syn-PFF-induced ser129 phosphorylation of α-syn S129 and α-syn accumulation. **A,** Western blot assay of α-synuclein pre-formed fibril (α-syn-PFF) based on human α-syn antibody (n = 3 per group). **B,** α-syn-PFF was stained with Coomassie blue. **C,** Transmission electron microscopy results of recombinant human α-syn-PFF at 10,000×magnification. **D,** The expression level of α7-nAChR in the PD (Parkinson’s disease) phenotype-like model was assessed using western blotting after feeding nicotine-containing drinking water for 1 month (n = 3 per group). **E,** Immunofluorescence staining was performed to detect the expression of the phosphorylated form of α-syn ser129 (green). **F,** The expression of α-syn aggregates (green). **G,** The expression of the α-syn aggregation marker p62 (green) in the hippocampus. \**P* < 0.05; \*\**P* < 0.01.

The α7 nicotinic acetylcholine receptor (α7-nAChR) is an important nicotine receptor (Saw et al. 2021) and has abundant α7-nAChR expression in the mouse hippocampus (Hogg et al. 2003). The antibody to α7-nAChR was used to detect nicotine intake in mice, and the results showed a significant upregulation of α7-nAChR expression in the hippocampus of mice fed nicotine-containing drinking water, with more significant α7-nAChR activation in mice injected with α-syn-PFF + nicotine group (Fig. 1D).

Approximately 90% of the α-syn protein deposited in the Lewy bodies of PD patients is phosphorylated at serine 129 (Ser129), suggesting that post-translational modifications of α-syn, particularly phosphorylation, are associated with the pathogenesis of PD (Anderson et al. 2006; Kam et al. 2018). Similarly, we detected α-syn serine 129 phosphorylation in the hippocampal region of mice injected with α-syn-PFF (Fig. 1E) and α-syn aggregates (Fig. 1F). The protein p62 is a ubiquitin-binding protein encoded by Sequestosome-1 (an important selective autophagy junction protein) and is a selective substrate of autophagy, the presence of which is frequently detected in abnormal protein aggregates and inclusion bodies formed in cells (Ichimura and Komatsu 2010; Su and Wang 2011). We also detected p62 accumulation in the hippocampus of the constructed α-syn-PFF mouse model (Fig. 1G). Simultaneously, the expression levels of α-syn Ser129 in the phosphorylated form, aggregates, and p62 were reduced in the hippocampus of mice fed nicotine-containing drinking water (Fig. 1E–G), and no significant accumulation of these proteins occurred in the hippocampus of control mice injected with sterile PBS and fed nicotine-containing drinking water.

### Nicotine improves *α*-syn-PFF-induced motor deficits, cognitive impairment, and anxiety-like behavior

Previous studies have found that injection of mouse α-syn-PFF into the hippocampus of mice leads to cognitive impairment (Nouraei et al. 2018). The mice in our model showed significant behavioral changes along with the accumulation of α-syn pathology. Mice with similar motor abilities (Fig. 2A) were randomly grouped, and 1 month after human α-syn-PFF injection into the hippocampus, the number of times the mice entered the novel arm, as well as the distance and time spent moving in the novel arm were significantly reduced (Fig. 2B), indicating impaired learning and memory. Moreover, the total distance and average speed of spontaneous activity in the open field (Fig. 2C,D) were significantly reduced, and the time spent on the spinning bar was also decreasing(Fig. 2E), indicating that the locomotor abilities of the mice were diminished. We also noted a significant decrease in the number of times the mice entered the central area of the open field by the open arm of the O maze (Fig. 2F,G,H), indicating anxiety-like behavior. Furthermore, when mice injected with α-syn-PFF were fed nicotine-containing drinking water for 1 month, memory and motor deficits, as well as anxiety-like behavior improved (Fig. 2B-H). Additionally, mice injected with sterile PBS and fed nicotine-containing drinking water showed impaired learning and memory, but this did not affect their motor performance or anxiety.

**Fig. 2.**
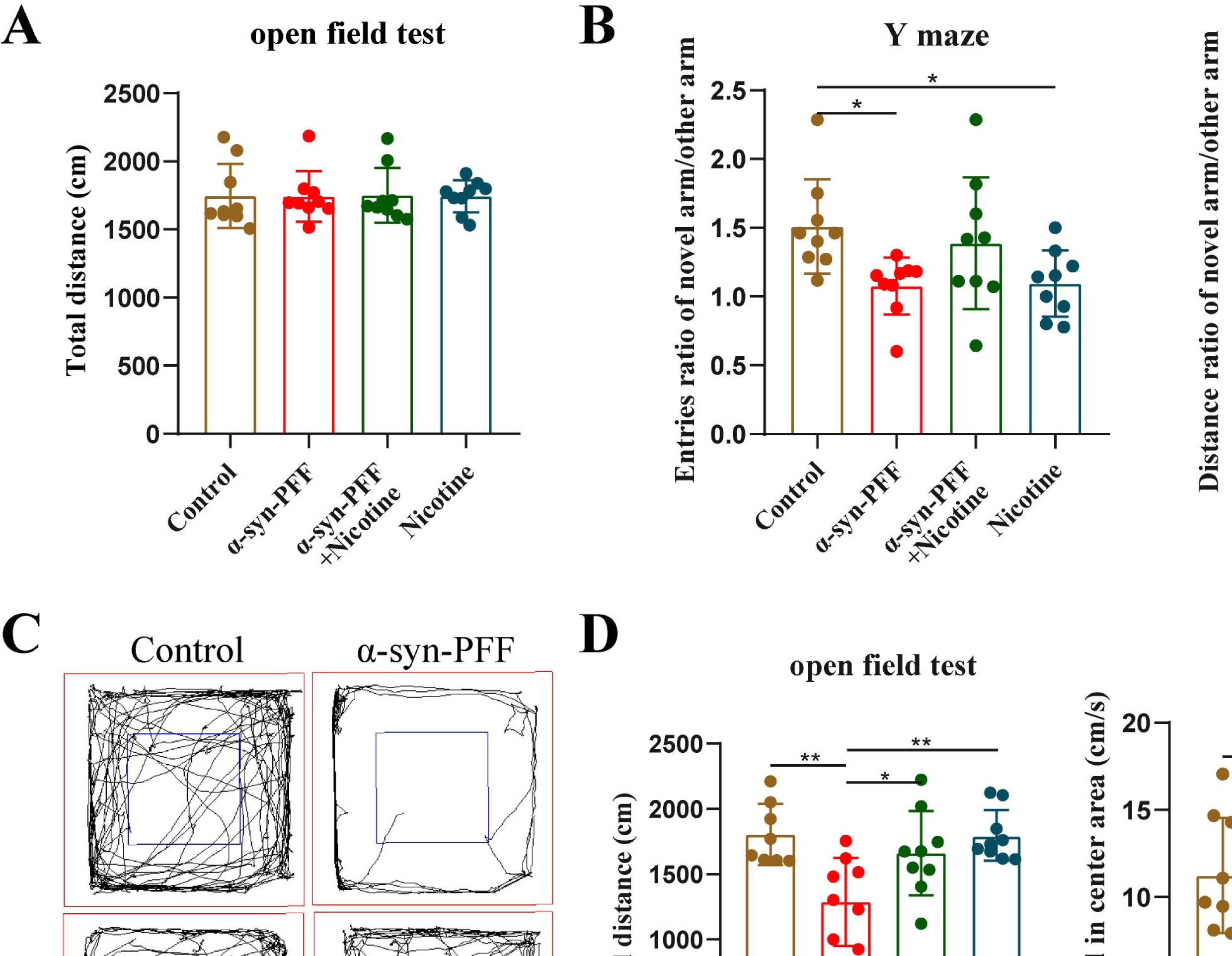
Mouse model constructed by hippocampal injection of α-syn-PFF exhibited significant memory impairment and motor deficits that were somewhat ameliorated by nicotine. **A,** Statistical graph showing the spontaneous movements of all mice in the open field experiment before performing stereotaxic injection surgery and before nicotine administration (n = 10 per group). **B,** Statistical graphs showing the ratio of the number of times that the mice entered the novel arm to the other arms in the Y maze, the ratio of the distance traveled in the novel arm, and the ratio of the time spent in the novel arm (n = 10 per group). **C,** A plot of the trajectory of mice in the open field. **D,** Statistical graphs showing the distance traveled and central and mean peripheral velocity of mice moving in the open field (n = 10 per group). **E,** Statistical graph showing the length of time spent turning the stick (n = 10 per group). **F,** A plot of the trajectory of mice in the O maze. **G,** Statistical graphs of the number of times that mice entered the central area of the open field (n = 10 per group). **H,** Statistical graph of the number of times that mice entered the open arm of the O maze, and the time spent in the open and closed arms (n = 10 per group). \**P* < 0.05, \*\**P* < 0.01, \*\*\**P* < 0.001, and \*\*\*\**P* < 0.0001.

### Nicotine inhibits *α*-syn-PFF-induced glial cell proliferation

The occurrence of PD is closely related to neuroinflammation. Therefore, we next used immunofluorescence and western blotting to detect the number and the expression of glial cells in the hippocampus after injection of α-syn-PFF, and the results showed that both microglia (Iba1) (Fig. 3A) and astrocytes (GFAP) (Fig. 3B) had proliferated, and the expression of Iba1 and GFAP were upregulated most significantly in α-syn-PFF group (Fig. 3C). However, nicotine significantly reduced the number of activated microglia (Fig. 3A) and astrocytes (Fig. 3B), at the same time, the expression of Iba1 and GFAP was also reduced (Fig. 3C).

**Fig. 3.**
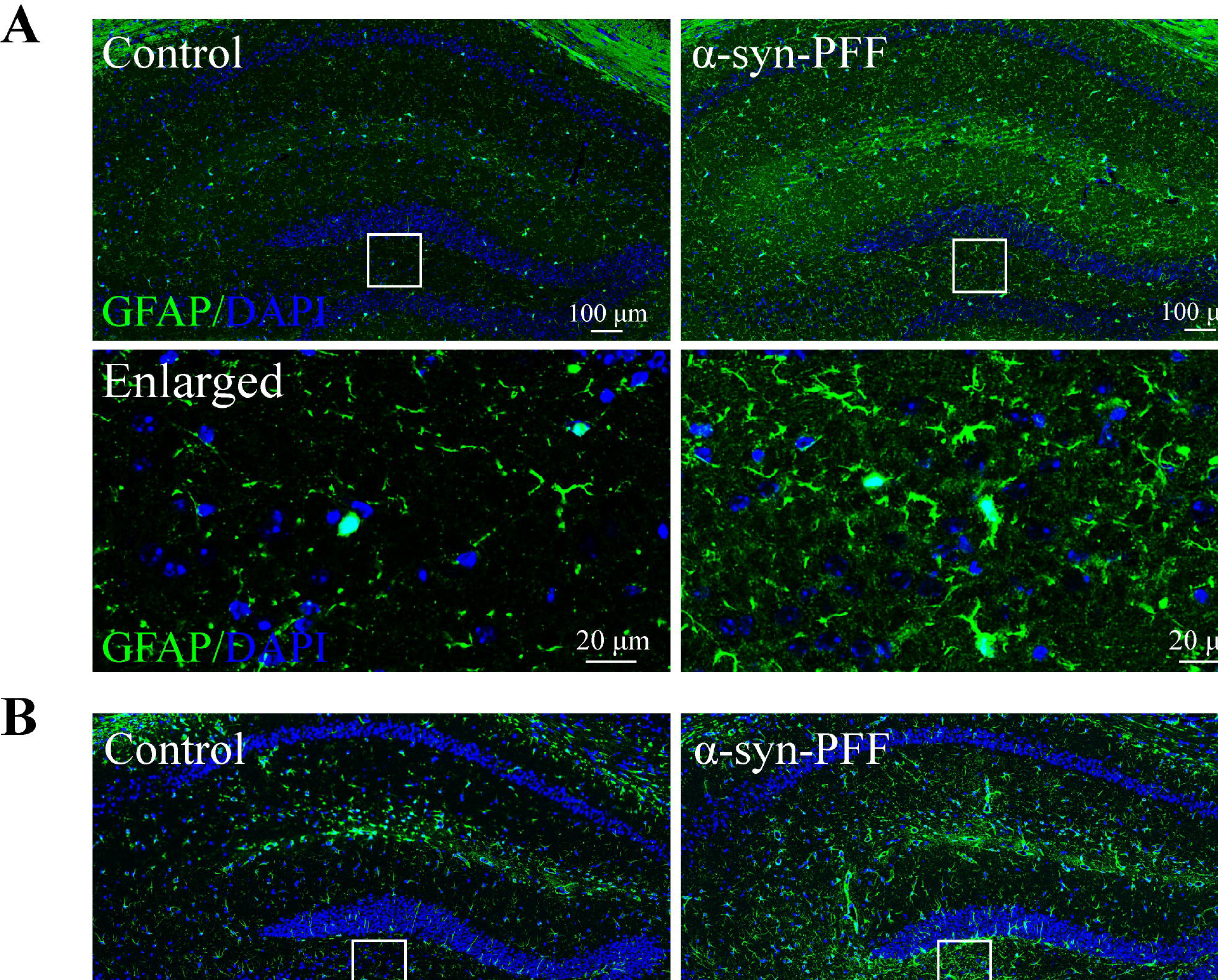
Nicotine reduces α-syn-PFF-induced glial cell proliferation. **A-B,** Immunofluorescence labeling of the mouse hippocampus with Iba1 (green) and GFAP (green) antibody at 1 month after α-syn-PFF injection; DAPI was used to stain all nuclei (blue). **C,** Western blot and statistical plots of the protein levels (ratio to GAPDH) with Iba1 and GFAP antibodies (n = 3 per group) in the mouse hippocampus. \*\**P* < 0.01; \*\*\**P* < 0.001.

### Nicotine inhibits *α*-syn-PFF-induced apoptosis

Activation of microglia and astrocytes results in the production of an excess of pro-inflammatory factors, inducing a severe chronic inflammatory response, which can result in neurological tissue damage. The proteins p53, Bax, and C-Cas3 involved in mediating apoptosis were also significantly upregulated in the hippocampus (Fig. 4A). The immunofluorescence results showed a significant increase in the expression of the microglial phagocytic marker CD68 after injection of α-syn-PFF (Fig. 4B). Both C-Cas3 immunofluorescence and TUNEL staining revealed an abundance of apoptotic cells in the hippocampus of mice injected with α-syn-PFF, especially in the CA1 and CA2 regions (Fig. 4C,D). The number of apoptotic cells in the hippocampus was significantly reduced after 1 month of feeding nicotine-containing drinking water to the mice injected with α-syn-PFF (Fig. 4A–D). Meanwhile, mice injected with sterile PBS and fed nicotine-containing drinking water showed only a small upregulation of CD68 in the hippocampus (Fig. 4B), with no significant apoptosis (Fig. 4 A,C,D). Nicotine intake activated the phosphatidylinositol 3-kinase-protein kinase B (PI3K-Akt) signaling pathway in mice in contrast to the α-syn-PFF group, and a significant upregulation of PI3K and Akt protein expression was detected in the hippocampus of mice fed nicotine-containing drinking water using western blot assays (Fig. 4E).

**Fig. 4.**
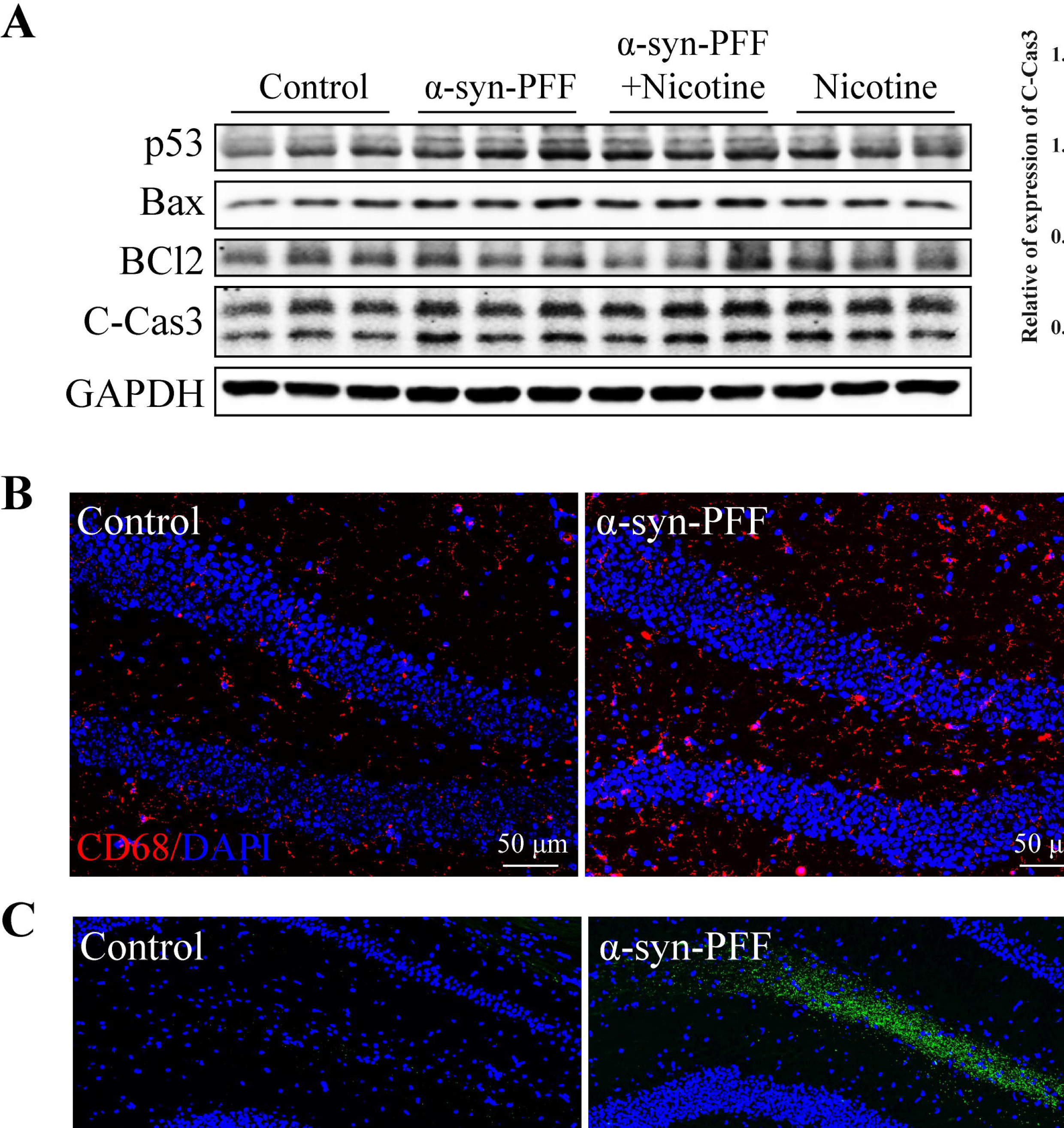
Nicotine inhibits α-syn-PFF-induced apoptosis of neurons in mouse hippocampus. **A,** Western blot and statistical plots of protein levels (ratio to GAPDH) with antibodies to p53, Bax, BCl2, and C-Cas3 (n = 3 per group) in the mouse hippocampal neuron. **B-C,** Immunofluorescence labeling of the mouse hippocampus with the microglia phagocytic marker CD68 (red) and cleaved-caspase3 (green) antibodies 1 month after injection of α-syn-PFF; DAPI was used to stain all cell nuclei (blue). **D,** The hippocampal region was stained for TUNEL (red) (n = 3 per group). **E,** Western blot and a statistical plot showing the protein levels (ratio to GAPDH) with PI3K and Akt antibodies in the mouse hippocampus (n = 3 per group). \**P* < 0.05, \*\**P* < 0.01, and \*\*\**P* < 0.001.

### Nicotine inhibits *α*-syn-PFF-induced neural stem cell reduction

The dentate gyrus is rich in neural stem cells, and the number of neural stem cells in the hippocampus was detected using antibodies to the sex-determining region Y-Box2 (SOX2) and doublecortin (DCX). The immunofluorescence results showed that the numbers of both SOX2 and DCX-positive cells in the dentate gyrus of the mouse hippocampus decreased after α-syn-PFF injection (Fig. 5A, B). Additionally, the apical spines of dendrites of neural stem cells labeled with DCX antibody in the hippocampus of control mice extended into the molecular layer, while the dendritic bifurcation and spine length of DCX-labeled neural stem cells were significantly reduced following injection of α-syn-PFF (Fig. 5C). The reduction in cell number and the morphological changes of neural stem cells were somewhat inhibited by feeding with nicotine-containing drinking water. Simultaneously, control mice fed nicotine-containing water showed no significant changes in the number or morphology of neural stem cells. Thus, nicotine could improve the reduction in the number and morphological changes of neural stem cells induced by α-syn-PFF. We quantified the number of SOX2^+^/DAPI^+^ and DCX^+^/DAPI^+^ cells in the hippocampal dentate gyrus at corresponding levels in a subset of mice (n = 3) from each group (Fig. 5D, E). The labeled cells were counted in eight randomly selected regions by blind observers at 40-fold target values. Green fluorescent and DAPI double-labeled cells were SOX2 and DCX-positive cells, respectively. Subsequently, the percentage of cells positively expressing each antibody was calculated and averaged.

**Fig. 5.**
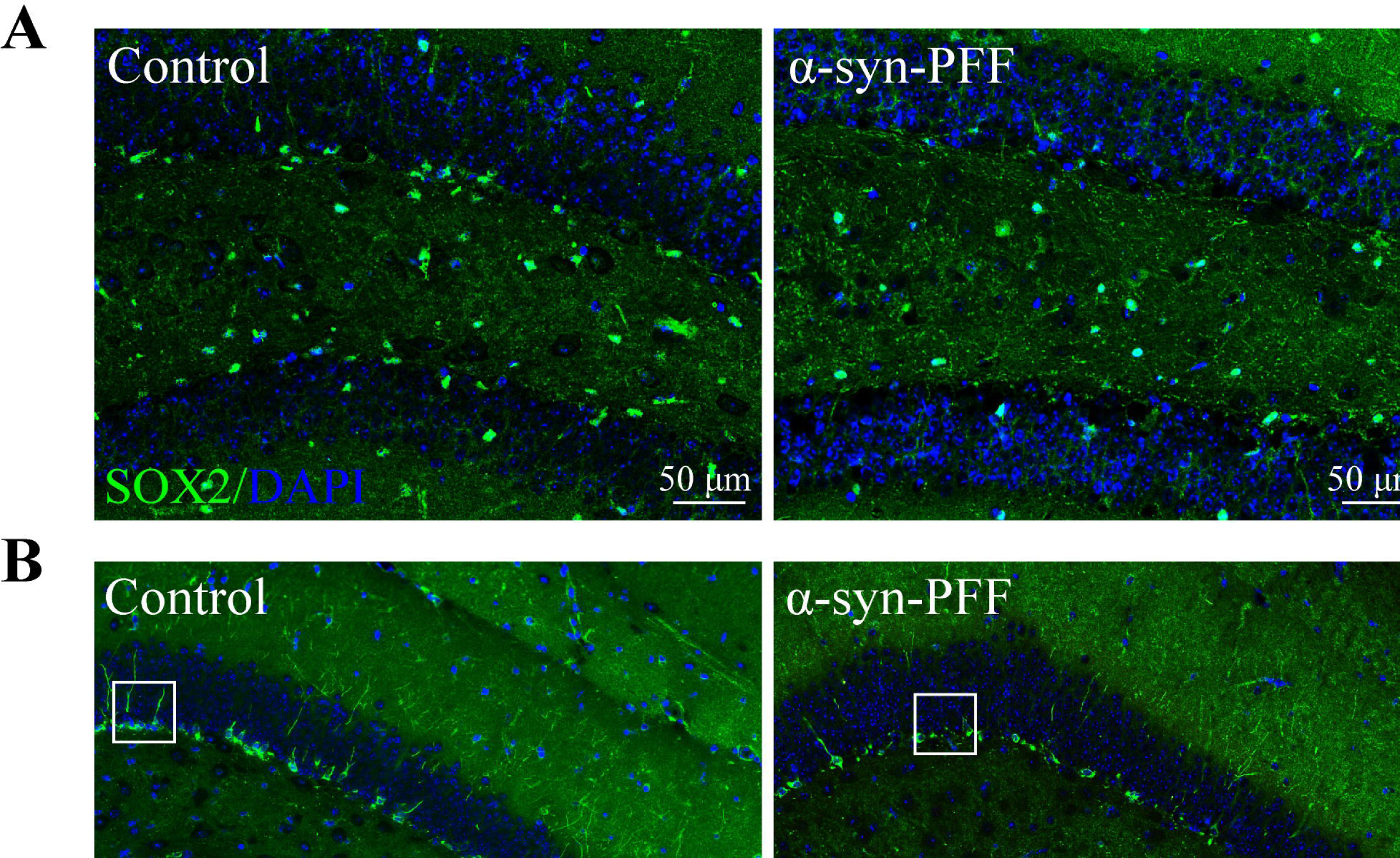
Nicotine inhibits α-syn-PFF-induced neural stem cell reduction. **A-B,** The expression of sex-determining region Y-Box2 (SOX2) and doublecortin (DCX) was detected using immunofluorescence 1 month after injection of α-syn-PFF; DAPI was used to stain all nuclei (blue). **C,** Morphology of DCX-positive cells in each group at 1 month after α-syn-PFF injection; DAPI was used to stain all nuclei (blue). **D,** Representative SOX2 and DCX-positive cells staining image at month 1 after α-syn-PFF injection. **E,** Quantification of SOX2 and DCX-positive cells at months 1 after α-syn-PFF injection (n = 4 per group). \*\**P* < 0.01, \*\*\**P* < 0.001, and \*\*\*\**P* < 0.0001.

## Discussion

Previous studies have found that injection of murine-derived α-syn-PFF into the striatum or hippocampus of the mouse brain induced the production of large numbers of LB-like inclusion bodies and that inoculation of α-syn-PFF in primary neuronal cells induced the production of large numbers of inclusion bodies, with a significant increase in the level of serine phosphorylation at position 129 of α-syn (Kam et al. 2018; Zhang et al. 2019). Our findings were similar. Compared with the classical toxin model and transgenic model of PD, the α-syn-PFF-based animal models of PD feature induction of the production of endogenous α-syn at levels that are closer to human physiological levels (Polinski et al. 2018). Thus, PD-related pathological models based on α-syn fibrils have unique advantages. Previous results have shown that inoculation of synthetic α-syn fibrils in the mouse striatum results in the intercellular spread of pathological α-syn and Parkinson’s-like Lewy’s pathology in anatomically interrelated areas, with reduced motor coordination but no significant impact on working memory, even 6 months after inoculation, and that mice did not develop pathology in the hippocampus (Luk et al. 2012). However, it is possible that 6 months after α-syn-PFF treatment was insufficient for the protein to spread from the striatum to the hippocampus. In this study, α-syn-PFF was injected locally in the hippocampus of mice; 1 month later, we observed a significant accumulation of α-syn in the hippocampal region, which resulted in neuroinflammation and the induction of behavioral deficits in mice, including mood disorders, memory impairment, and deficits in motor behavior. Given the advantage of rapid production of abnormal behavioral and pathological features in the model coupled with the administration of nicotine at a low dose for 1 month, this modeling method partially reduced cognitive deficits in novel object recognition and social impairment.

Nicotine administered orally (dissolved in tap water) has been used in various experiments, including studies of addiction, withdrawal, and toxicity to offspring (Grabus et al. 2005; Matta et al. 2007; Zhao-Shea et al. 2015; Mojica et al. 2018). Therefore, we injected α-syn-PFF into the hippocampus of mice to induce rapid abnormal behavior, while the test group was fed nicotine-containing drinking water (200 µg/mL) for 1 month. Current research has shown that nicotine can, to some extent, reduce the α-syn phosphorylation of serine 129 induced rapidly by α-syn-PFF, as well as inhibit the aggregation of α-syn. This is consistent with the results obtained by Zhao et al., who demonstrated through SH-SY5Y cell assays and knockout mouse models that the activation of α7-nAChRs had a protective effect against α-syn-PFF-induced and mitochondrial apoptosis and that this was achieved by inhibiting the accumulation and phosphorylation of α-syn, ultimately leading to neuroprotection in PD mice (Zhao et al. 2021). Our results also indicate that a certain dose of nicotine produces a rapid response against hippocampal damage because nicotine can bind to α-syn-PFF and inhibit α-syn aggregation by reducing α-syn phosphorylation at the Ser129 position.

One of the ways in which α-syn accumulates to produce oxidative stress is through the activation of microglia, and microglia-derived factors, such as pro- and anti-inflammatory factors and oxidative stress substances, can trigger astrocyte receptors, thereby inducing inflammation that can cause neurological damage (Jha et al. 2019). Nicotine is a cholinergic receptor agonist that mimics the binding and activation of acetylcholine to nAChRs (Karban and Eliakim 2007). Moreover, it has been reported (van Westerloo et al. 2005) that nicotine inhibits microglia activation and thus protects dopaminergic neurons. Indeed, in an MPTP mouse model of PD, the systemic application of nicotine resulted in the upregulation of α7-nAChR expression and counteracted the loss of dopaminergic neurons and the activation of astrocytes and microglia (Liu et al. 2012). The results of in vitro experiments also demonstrated that α7-nAChR in microglia is a key regulator of the peripheral inflammatory response (Wang et al. 2003). Using lipopolysaccharide (LPS)-induced inflammation models in vitro and in vivo, a previous research demonstrated that nicotine exerts anti-inflammatory effects by reducing the activation of microglia and astrocytes and significantly decreasing the expression and release of endotoxin stimulation-induced tumor necrosis factor-α (Park et al. 2007). In the present study, compared with mice in the α-syn-PFF injection group, those in the α-syn-PFF + nicotine group had significantly reduced expression of both the microglia marker protein Iba1 and the astrocyte marker protein GFAP in the hippocampus. Our results suggest that nicotine prevents the neuroinflammatory response induced by α-syn-PFF in the hippocampus of mice. Previous studies have shown that the reduction in neurogenesis is one of the consequences of neuroinflammation (Gebara et al. 2016) and that excess pro-inflammatory factors may contribute to the quiescence of more neural stem cells and lead to the failure of the stem cell pool. Here, we observed a similar phenomenon, where the activation of microglia and astrocytes caused damage to mouse hippocampal neurons. Additionally, the protein expression levels of p53, Bax, and C-Cas3, which are involved in mediating apoptosis, were significantly upregulated, and apoptosis was increased in the hippocampus of mice injected with α-syn-PFF compared with the control group. However, these effects of α-syn-PFF on the hippocampus of mice were reversed by nicotine treatment. The treatment inhibited the expression levels of the microglia phagocytosis marker CD68, promoted synaptogenesis in the hippocampus, led to increased expression levels of the neural stem cell markers DCX and SOX2, and increased the number of DCX+ cells, whereas mice fed only nicotine-containing drinking water showed no significant changes in the number of neural stem cells. In vivo studies have shown that chronic administration of nicotine with periodic deprivation increases the number of immature neurons in the hippocampal region (Cohen et al. 2015b). Nicotine also inhibits dopamine neural toxicity in mice caused by rotenone through the activation of nAChR (Takeuchi et al. 2009). Nicotine has also been suggested to protect dopaminergic neurons from apoptosis by reducing endoplasmic reticulum stress and unfolded protein responses (Srinivasan et al. 2016). Therefore, we conclude that nicotine protects against α-syn-PFF-induced apoptosis and inhibits α-syn-PFF-induced neural stem cell reduction by inhibiting α-syn aggregation and upregulating α7-nAChR expression. This is also reflected by the findings of recent studies, which have shown that both the α7-nAChR agonist PNU-282987 and nicotine reduced apoptosis induced by α-syn-PFF and that α7-nAChR activation inhibited the expression of the apoptotic proteins Bax and C-Cas3 and maintained mitochondrial membrane potential in a mouse model of PD (Zhao et al. 2021).

Nicotine activates the PI3K-Akt signaling pathway via α7 nAChR (Kawamata et al. 2012), and the activation of Akt increases the expression of the apoptosis regulatory gene B lymphocytoma-2 protein (BCl2), a negative regulator of apoptosis that protects cells from apoptosis and promotes p53 protein degradation to facilitate survival. Our results demonstrate that PI3K and Akt protein expression was significantly upregulated in the hippocampi of mice after nicotine intake compared with those in the α-syn-PFF group, thereby activating the PI3K-Akt signaling pathway, inhibiting apoptosis, and protecting neural stem cell survival. The increase in neural stem cells and new neurons in the hippocampus, a key brain region responsible for learning and memory, plays an important role in improving cognitive function and coping with stress and anxiety in mice. Furthermore, a decrease in neural stem cells may lead to a decline in hippocampal neurogenesis, thus affecting the normal function of the hippocampus and leading to memory impairment and anxiety-like behavior (Zhang et al. 2008). The same behavioral manifestations, along with motor deficits, were observed in the present study in mice in which α-syn-PFF was injected into the hippocampus, while nicotine intake resulted in significant improvements in these behaviors. Additionally, there is evidence that nAChR is significantly reduced in the post-mortem cerebral cortex of patients with PD with dementia and non-dementia (Everitt and Robbins 1997; Oishi et al. 2007). Recently, researchers knocked out nicotinic receptors from astrocytes and found a significant reduction in fear memory behavior (Zhang et al. 2021). Moreover, nicotine improved memory and selective attention deficits and consolidated functional thalamocortical connectivity in patients with schizophrenia (SCZ) by stimulating α7-nAChR (Mehta et al. 2019). Nicotine upregulated α7-nAChR expression and activated astrocytes and microglia while reducing MPTP-induced behavioral symptoms and improving motor coordination in mice (Liu et al. 2012). As demonstrated by the rotenone rat model of PD, nicotine partially blocked the reduction of dopamine levels in the striatum and ameliorated motor deficits (Mouhape et al. 2019). In non-human primates, nicotine reduced levodopa-induced dyskinesia in rhesus macaques (Quik et al. 2007; Quik et al. 2013). Further studies have shown that α4β2* and α6β2* nAChRs play an important role in the anti-motor effects of nicotine (Huang et al. 2011b). Our results appear to better explain the ability of nicotine to alleviate the α-syn-PFF-induced non-motor symptoms, such as anxiety and cognitive impairment, in addition to the motor symptoms caused by the hippocampal injection of α-syn-PFF. Although a previous study suggested that nicotine significantly improved cognitive impairment caused by overexpression of α-syn, nicotine did not alter α-syn aggregation and did not inhibit astrocyte activation (Subramaniam et al. 2018). This was not entirely consistent with our findings, although the discrepancy may be explained by the different dose or route of administration. Indeed, researchers have suggested that the mode of administration is an important consideration when using nicotine because nicotine receptors are rapidly desensitized with continued exposure to nicotine, as in the case of the nylon patch (Giniatullin et al. 2005; Dani and Bertrand 2007). Previous studies have confirmed the ability of nicotine to exert broad protective effects, which may be important in PD, where neuronal deficits are known to extend throughout the peripheral and central nervous system (Braak et al. 2003; Braak et al. 2006); this observation seems to better explain our findings.

Although the present study shows that nicotine plays a role in improving the behavior and hippocampal pathology of α-syn-PFF-induced mice, its deficiencies cannot be ignored. Through the O maze and Y maze tests, we found that control mice of the nicotine group also showed deficits in cognitive learning, as well as elevated expression of microglial markers (Iba1) and astrocytic markers (GFAP) in the hippocampus. A similar study recently reported that chronic exposure to nicotine caused decreased hippocampal neurogenesis in mice, resulting in slow development and impaired memory (Nakayama et al. 2019). Maternal nicotine exposure also affected hippocampal neurogenesis and microglial hyperactivity in the offspring, with activated microglia increasing the release of various pro-inflammatory cytokines and promoting anxiety and depression-like behavior. This suggests that nicotine contributes to cognitive impairment through other mechanisms and pathways, highlighting the need to consider both the duration and cumulative amount of nicotine used in the context of PD. Further research is needed on the paradox between the positive and negative effects of nicotine treatment.

## Conclusions

In the current study, we rapidly induced cognitive deficits, motor deficits, and anxiety-like behavior in mice by stereotactic injection of α-syn-PFF in the hippocampus, which resulted in significant pathological accumulation of α-syn in the hippocampus. However, the administration of nicotine stimulated α7-nAChR to inhibit the pathological accumulation of α-syn and maintained neurogenesis by reducing hippocampal neuronal apoptosis, possibly through inhibition of glial cell proliferation and activation of the PI3K-Akt signaling pathway, thereby reducing behavioral deficits induced by α-syn fibrils. The results suggest a role for nicotine in α-syn-PFF-induced behavioral deficits and hippocampal pathological abnormalities in mice, indicating a potential downstream molecular mechanism for the neuroprotective effects of nicotine in PD in Fig. 6. However, the decline of cognitive and motor ability involves damage to hippocampal and cortical circuits; whether α-syn fibrils spread to the cortex and other brain regions and whether nicotine can inhibit the spread of α-syn fibrils require further study.

**Fig. 6.**
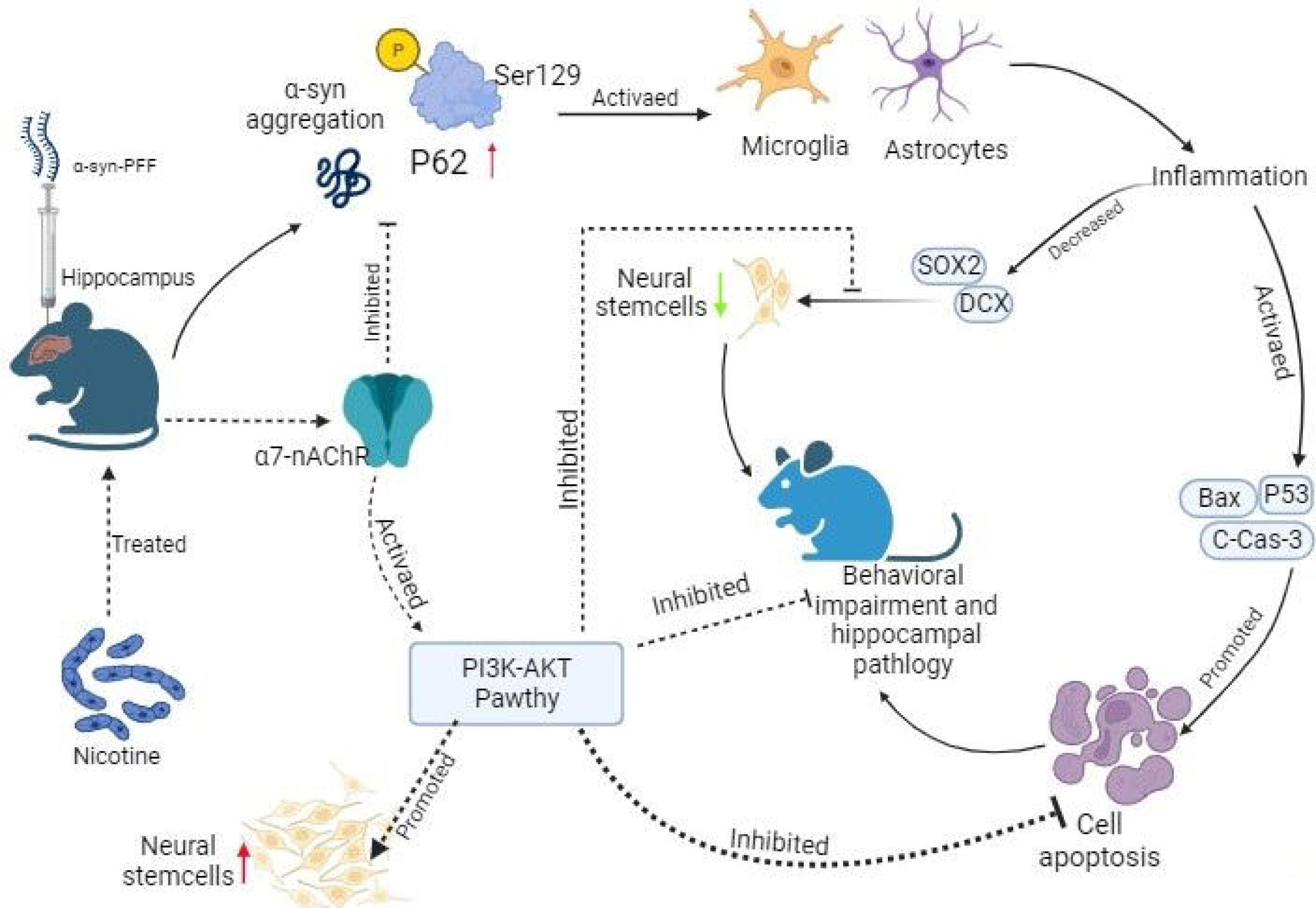
Schematic illustrating the nicotine-induced amelioration of α-synuclein fibril-induced behavioral deficits and pathological features in mice. A significant pathologic accumulation and phosphorylation at the Ser129 position of α-syn occurred in the hippocampus 1 month after α-syn-PFF injection, and this activated microglia and astrocytes, leading to inflammation, apoptosis, and a reduction in neural stem cells, ultimately resulting in behavioral impairment and hippocampal pathology in mice. However, after the administration of nicotine, α7-nAChR could be activated to inhibit the pathological accumulation of α-syn and maintain neurogenesis by reducing hippocampal neuronal apoptosis, concurrently, through the activation of the PI3K-Akt signaling pathway, ameliorating behavioral deficits induced by α-syn fibrils. The schematic illustration was drawn using BioRender (https://biorender.com/).

## Abbreviations

PD: Parkinson’s disease
α-syn: Alpha-synuclein
LBD: Lewy body dementia
RIPA: Radio Immunoprecipitation Assay
PMSF: Phenylmethanesulfonyl fluoride
EDTA: Ethylene Diamine Tetraacetie Acid
DAPI: 4’,6-diamidino-2-phenylindole
TUNEL: YF®594-dUTP notch nick end labeling
p62: Sequestosome-1(SQSTM1)
SQSTM1: Sequestosome-1
S129: Serine-129
C-Cas3: Cleaved-Caspase3
CA1: Hippocampal ca 1 region
CA2: Hippocampal ca 2 region
SOX2: Sex-determining region Y-Box2
DCX: Doublecortin
PBS: Phosphate buffer saline
WB: Western blot
IF: Immunofluorescence
CON: Control
NIC: Nicotine
α7-nAChR: α7 nicotinic acetylcholine receptor
α-syn-PFF: α-synuclein pre-formed fibrils
Iba1: Ionized calcium-binding adapter molecule 1
GFAP: Glial fibrillary acidic protein
PI3K: Phosphatidylinositol 3 kinase
Akt: Protein kinase B
SCZ: Schizophrenia
MPTP: 1-Methyl-4-Phenyl-1,2,3,6-Tetrahydropyridine (neurotoxin)
BCl2: B lymphocytoma-2 protein
GST: Glutathione S-transferase
AP: Anteroposterior
ML: Mediolateral
DV: Dorsoventral

## Acknowledgments

Thanks to LetPub Editing Services for doing a lot of research on the grammar and words of the article.

## Author‘s’ Contributions

G. Y. and Z.Z.H. proposed the study design. H. Z. Q. and P. Y. performed the additional experiments for insufficient data and discussion, and wrote the manuscript. L.H.Y. and Z. Q.L. performed the behavioral experiments and discussion. M.K. L. and W.Z.C. performed the statistical analysis and discussion. All authors reviewed the manuscript.

## Data Availability

The data used in the article is presented in the article/supplementary material. Please contact the corresponding author for further information.

## Declarations

Ethics approval and consent to participate

Ethical approval was obtained from the Experimental Animal Ethics Committee of the Institute of Medical Biology Chinese Academy of Medical Sciences (DWSP202203024). All methods were carried out in accordance with relevant guidelines and regulations.

## Consent for publication

Not applicable.

